# Multi-BIRW: a network-based algorithm for drug repurposing

**DOI:** 10.1101/2020.09.04.281451

**Authors:** Luigi Ferraro

## Abstract

This paper is based on *Computational Biology* with an eye towards medicine. In fact our aim is to identify new indications for existing drugs with a computational approach using *Systems Biology*, in order to speed up the market entry, avoiding some phases of drug development (for example toxicity tests) and with a lower cost.

The method that we will discuss exploits the concept of *Random Walk* in a heterogeneous network, formed by a drug-drug and disease-disease similarity and known drug-disease associations. This is done by combining different types of data in order to connect more effectively two drugs or two diseases with a similarity score. At the end we integrate the known associations between drugs and diseases with the purpose of finding similarity values of new couples.

## Introduction

Nowadays *Drug repurposing* is given more importance, i.e. the discovery of other indications for existing drugs, bearing in mind the significantly advantages in the exploitation of the process on behalf of the pharmaceutical companies. For example: the risk of failure is lower, less time-consuming and less investment is needed since many tests are already satisfied by the candidates.

Recently, many computational methods, to find other uses for a drug, are being developed on the grounds as they accelerate the process of discovering new associations. Considering all computational methods, one stands out above all: *Network based algorithm*. It could be used in order to combine different data source in a heterogeneous network. We will give some examples of current computational approaches.

Chiang and Butte [2] predict new association exploiting the assumption that if two diseases share similar therapies, subsequently drug used only for one disease could be considered as a new association with the other disease.

Wu et al. [12] use a weighted heterogeneous network connecting drug and disease which is constructed using known disease–gene and drug–target relationships and, in order to to find candidates, they applied a clustering operation.

Also Wang et al. [10] use a heterogeneous network. They apply drug repositioning and target prediction and, at the end of an iterative algorithm, candidates are sorted picking the top ones.

In this study we present a method based on two kind of application exploiting *Bi-Random Walk* and *Matrix Factorization* exposed respectively by [8] and [13]. These two insights were used efficiently in order to predict new indications for existing drugs, so our insight was to combine them in order to achieve better performance. An intuition of the workflow is given by Figure 1.

Given these basis, the diversity of these two methods leads us to a new perspective of the problem, which integrates more kind of data integrating different views. Therefore our results achieve better prediction respect to other methods, taking the best parts of the applications exposed before.

**Figure 1:**
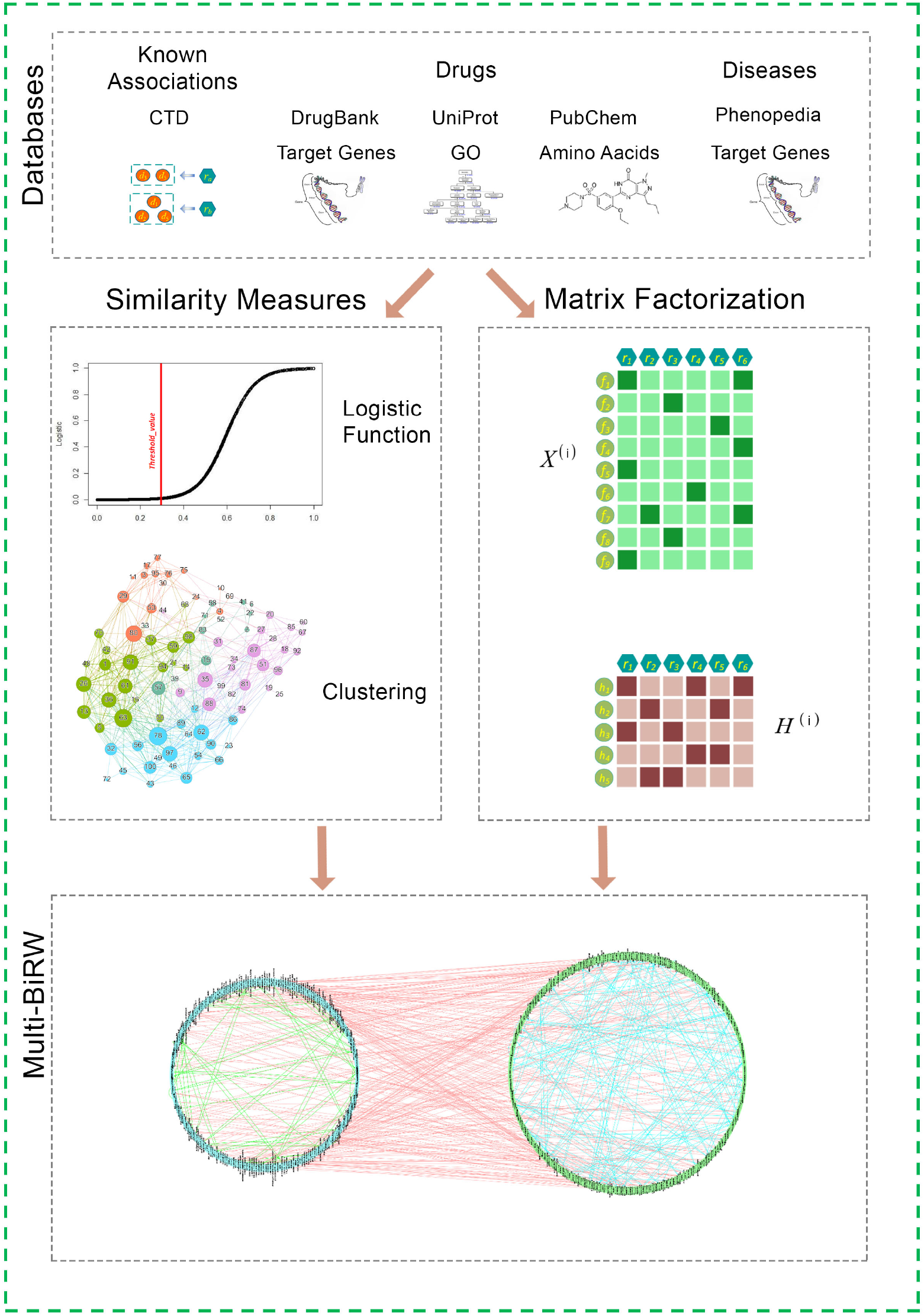
Workflow of our method

## Datasets

First of all, we had to choose what kind of data we want to manipulate.

Our initial intuition was that we required data that describe the known association between drugs and diseases, provided by *CTDatabase* [3]. Subsequently, the necessity to express the similarity between drug-drug and disease-disease led to finding of two databases *DrugBank* [11] for the drug side, *Phenopedia* [14] for disease side, which focus on this kind of interaction.

The correlation between diseases is sufficiently expressed by this kind of interaction, however, for the drug side the information is to be considered not satisfying. This led to the finding of other two databases that gather information about the targeted protein’s properties and what amino acids it is composed of. The first information is provided by *UniProt database* [4], the second by *PubChem database* [6].

### Manipolation of data

Data in *UniProt database*, *Phenopedia* and *DrugBank* are all represented by a field indicating which drug or disease they are and by another field containing the desired information (gene involved or property of targeted protein). We need to transform this kind of data in an appropriate matrix. First of all, we searched only for drugs and diseases which were in all databases. The matrices were attained considering the attributes of the columns in terms of drugs or diseases while the rows in terms of the genes involved or property of targeted proteins. The cells have a binary format, 1 if there is an association between drugs/diseases and the informations above mentioned, 0 otherwise. *PubChem database* provides a matrix already in this form.

## Similarity measures

First of all, the first method [8] modifies the similarity values according some statistical operation and at the end it makes uses of the clustering in order to improve them.

To indicate the similarity between two drugs (or two diseases) we have performed the matrix multiplication between the binary matrix with itself transposed. The resulting matrices indicates how many genes (property of targeted proteins, amino acids) they have in common.

The similarity is to be expressed as a value between 0 and 1, hence the following normalization was performed: the diagonal of the matrix, which indicates how many genes (property of targeted proteins, amino acids) have in common with itself (that is the maximum value of that row and that column), is multiplied by itself in order to get a matrix and finally the original matrix is divided by the squared root of the resulting matrix.

This kind of operation takes the name of **cosine similarity** and its formula is:

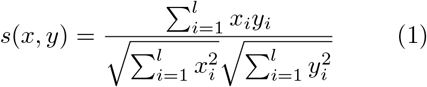

We used a similar approach, but totally equivalent and faster in computational time.

The *cosine similarity* acts like the *Tanimoto score* when the features are binary, as in our case. It is used to calculate the *Jaccard coefficient* when sets are viewed as bit vectors. Here is the method to calculate the *Tanimoto score*:

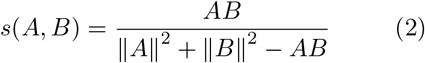

From now on we will refer to **Wdd** as the matrix that stores the similarity value between two diseases, to **Wrr**s as the list of matrices that stores the similarity values between two drugs and to **Wdr** as the binary matrix which express the known association between a disease and a drug. In particular, the similarity analyses are done for *Wdd* and for each matrix in *Wrrs*, so we will explain every step using only the alias **W**.

Inline with [8], a threshold is computed underneath which the similarity is to be considered not enough, so its value will be reduced. On the other hand those similarity values which are higher than the threshold will be rewarded.

In order to find communities in which drugs or diseases tend to behave similarly, we use clustering on the whole graph, perhaps leveraging the found similarities.

Our method is based on *Louvain clustering* [1]. It is a greedy algorithm created by Blondel and it revealed to be really efficient. In fact it runs in a time *O*(*nlog^2^n*) where *n* is the number of nodes in the graph. It is used for finding community in a weighted network (as in our case, since our graph is composed by edges representing the similarity between two nodes).

Subsequently the algorithm runs over the weighted adjacency matrix, we calculate a *quality* for each community as follow:

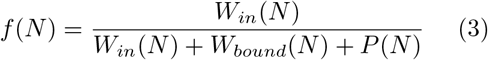

where:

- *W_in_*(*N*) is the number of edges within a group of nodes N;
- *W_bound_*(*N*) is the number of edges connecting this group to the rest of the graph;
- *P*(*N*) is a penalty term that is set as the percentage of nodes in the cluster.

This new similarity will update the current one of the nodes in this modality:

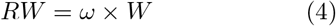

where *ω* is 1 + *f*(*N*). Some values could be more than 1, so we set them to 0.99. We also take the maximum value between the elements of *RW* and *f*(*N*) and store in *RW*. Subsequently the diagonal (corresponding to the similarity of each node with itself) is filled of 0s.

## Matrix factorization

The second method is based on the work of [13], which uses the concept of *Matrix factorization*.

The origin of this approach came in order to satisfy the problem arisen by Netflix [7]: “given a set of movies watched by user, which movie is better to recommend watching?”. The idea was to represent users and elements in an lower-dimensional matrix.

Our insight was to represent the drug-disease similarity using this idea, exploited by a non-negative matrix factorization.

The first aim of this approach is to decompose the drug features matrices into a matrix in lower dimension while was then given a shape in the form of (*m, n*) instead of the original (*f_i_, n*), where *m* is the number of diseases, *n* is the number of drugs and *f_i_* is the length of the drug feature. To these three new matrices are given the name of *H^v^* and to the original matrices the name of *X^v^*, where *v* is an index that ranges from 1 to 3, indicating which drug we are considering. We need a second type of matrices, which take the name of *W^v^*, representing the weight of each new features of *H^v^*, and has a shape of the form (*f_i_,m*).

According to this, the two terms of the next formula should be equal, or at least similar:

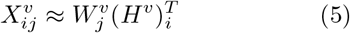

In line with [13]: “The multitude of drug similarities reflects the degree of similarity among the drugs from different aspects. There is consistency between the information from multiple aspects, but each view also has its own specific information”.

In order to keep the views of different features as various as possible, and so to be as orthogonal as possible, this paper introduces this new formula which should tend to zero:

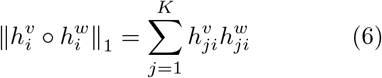

### Minimization problem

Our approach to this problem is to create a minimization problem which has to satisfy the two equations, see above Equation 5 and Equation 6. Using the Langrangian method, our constrained minimization problem is transformed in a unconstrained minimization problem. In order to do so, a objective function was created with the two constraints explained before.

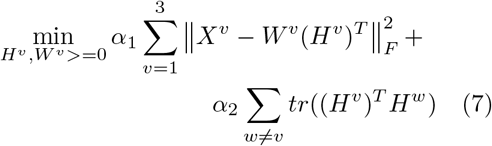

where 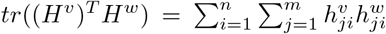 n is the number of drugs and m the number of diseases. The two alphas are factors of importance.

The aim is to find to find the values for matrices *H^v^* and *W^v^* that satisfy this minimization problem. That one may be solved using fixing values obtained from the loss function.

At each iteration, *H^v^* and *W^v^* are updated according to the following formulas, the value of loss function is calculated using Equation 7 and the new value is compared with the previous one: if their difference is less then the new loss value multiplied by a percentage (we set it to 0.005), the values are returned. The updating equations are found with the Langrangian method exploiting the fifth *KKT condition* [5].

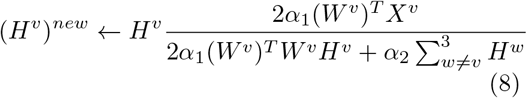

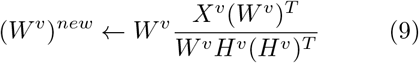

## BiRW

**Bi-Random Walk** is linked to the method used by [8]. It uses a *random walker* that exploits the heterogeneous network created by the combination of the two similarity matrices (*normWdd* and one of the list *normWrrs*) obtained in the final step of Figure and the matrix representing the known association between drugs and diseases *Wdr*, referred in Figure.

Given the mixed nature of the graph, we will refer to *left* and *right* part of random walker as the *drug* and *disease* side. This is the core point of the definition of the concept of **Bi-Random Walk**.

The steps of this tool are now explained.

First of all, we take the association matrix *Wdr* and we normalized it dividing *Wdr* by the sum of all matrix. The result is stored in two variables called *R*_0_ and *Rt*.

Following, for *n* times (in our experiments, *n* set to 2 seems to be solution), which represents the *walk steps* we applied this two formulas, for *left* and *right* side of the graph distinctively:

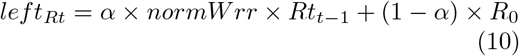

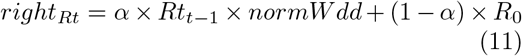

where:

- *α* is a *decay factor* set to 0.3;
- *Rt*_*t*−1_ is Rt at time *t* − 1, because it is updated at the end of every step, we will see how soon.

Here is what each term refers to:

- *Rt* in general stands for the new prediction matrix that links a drug and a disease with a certain probability;
- *normWrr* × *Rt*_t−1_ and *Rt*_*t*−1_ × *normW dd* are *updating terms*;
- *R*_0_ represents the *golden standard term*, this does not change at each iteration;
- *α* is important because gives a certain importance to the *updating terms*: a low *α* accentuates the *golden standard term*.

The random walker iteration could be done separately for *left* and *right* side. In fact *n* is the maximum value between *steps_left_* and *steps_right_* (set both to 2 as it revealed to be the the most valuable choice), which represent the number of iteration which each part must be taken into account.

At the end of the iteration, *Rt* is updated pooling *left_Rt_* and *right_Rt_*:

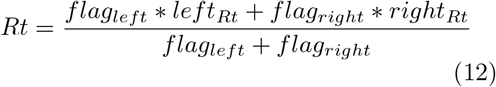

where *flag_left_* and *flag_right_* are Boolean value which indicates whether the iteration considers the *left* or the *right* side of the network, and are useful to calculate a weighted result when updating *Rt*.

### Multi Bi-Random Walk

Our insight was to introduce more Databases, and so more kind of data, for the drug sides. In fact our approach is similar to the first one, but we have three different graphs obtained combining *normWdd* with each matrix of the list *normWrrs*. In this way we have three Rt, three *left_Rt_* and three *right_Rt_*. Not only, the result of the *matrix factorization* are also considered, saved in the form of matrix in *H^v^*.

So the steps are like the first approach but in each iteration we get six different *Rt*, *left_Rt_* and *right_Rt_*, divided with two different methodologies: similarity measurements plus clustering and matrix factorization.

At the end of the iterations we consider these two kinds of *Rt* and separately we calculate a weighted sum of them divided by the number of features, getting two resulting matrices.

The resulting matrices may be in different order of magnitude, so we normalize them in this way:

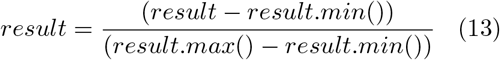

In this way both matrices, which we will call res**DP** and res**SIM**, have the maximum value equals to 1, and the lowest equals to 0.

At the end this we get the *top k* elements of each matrix, where *k* is equal to *n*\**m*\*1%, where *n* is the number of drugs and *m* is the number of diseases, and are then replaced in a score matrix (if the same association is considered by both, the maximum value is taken into account). Consequently, for each value that was not considered by the previous step, we take the maximum value between *resDP* and *resSIM*.

## Evaluation

In this chapter we will see how evaluation is performed in order to evaluate our algorithm. Then some statistics are then carried out about the combined network given in input to the Bi-Random Walk. Finally scientific basis are searched for new association.

To see how the prediction validity of our algorithm, we performed a *10-fold Cross-Validation*.

*K-fold Cross-Validation* is a method used to test a binary classification. First of all, we randomly split the *golden standard dataset* (known association between drugs and diseases) **Wdr** in *k* subsets (in our case 10). We cycled over this 10 groups, each time one of them become a *test set* while the remaining part of *Wdr* become *test set*. In order to do so, we set all the known associations of the test set equals to 0, so we get a new *Wdr*. Our aim is to find as many known associations of the test set as possible, which is assumed to be the most easy to discover.

The new *Wdr* is given as input to our algorithm, exhibited in the previous chapters. The results is, as always, a new *Rt*. From this, we take the *top k* association, where *k* this time refers to the size of test set, without considering the known association of the training set. Consequently the number of found associations, which where in the test set, are counted.

In order to give a graphical representation of our metrics, we used the **ROC curve** (*Receiver Operating Characteristic*) which plots the **True Positive Rate** (*TPR*) and **False Positive Rate** (*FPR*).

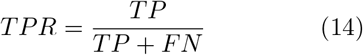

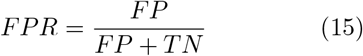

where:

- *TP* is the number of true positive, so the number of correctly found associations of test set;
- *FN* is the number of false negative, so the number of associations of test that were not found;
- *FP* is the number of false positive, so the number of found associations that were not in the test set;
- *TN* is the number of true negative, so the number of associations not found that were not in the test set.

Our *ROC curve* is rather satisfactory and it is shown in Figure 2. It considers the combination of truth tables of each iteration in the phase of *Cross-Validation*.

**Figure 2:**
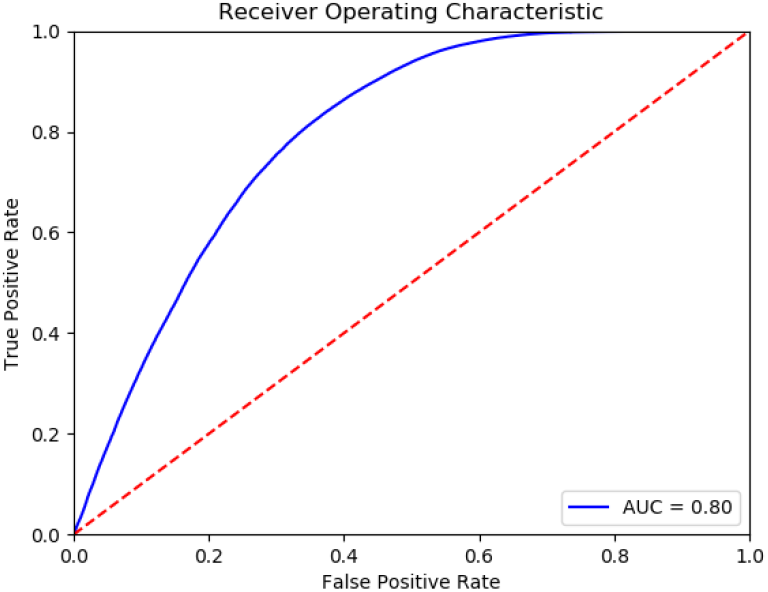
ROC curve

### Statistics of graph

Our graph is represented, in order to give a intuition of what we found.

Bearing in mind *Rt*, *normWdd* and one of *normWrrs* matrices, we used **Cytoscape** [9], which is an open-source program which performs some operation on large graphs and it helps with the visualization of them.

Our problem was that our graphs are weighted, so in the visualization part we would have seen a confusing graph with so many edges. Hence, to avoid this, we take the *top association* of each element. In case of *normWrrs*, first the diagonal is set to 0 (since the similarity of each drug with itself is not so explanatory), for each drug we keep the drug which is the most important for the starting one, similar for *normWdd* and diseases. In case of drug-disease association in *Rt*, the known relations are removed and considering the most important disease associated with each drug.

The first two elements with most edges are removed, since, from a graphical point of view, some nodes have numerous edges resulting in a confused figure.

With *Cytoscape* we combined the graphs of *normWdd*, *normWrrs* and *Rt*, as shown in Figure 3.

**Figure 3:**
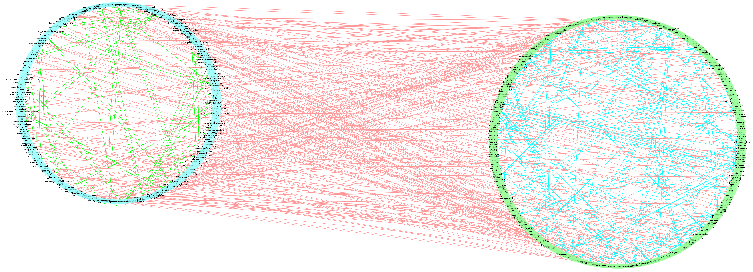
Merging three graphs: the one at the top is the similarity graph of diseases, the one at the bottom is the similarity graph of drugs, the red edges represent the association between a drug and a disease. The most important connection are chosen.

Subsequently some statistics have been performed about the resulting graph, some of them are shown in Figure 4 in form of *boxplots*, others are described as follows:in Table 1.

**Figure 4:**
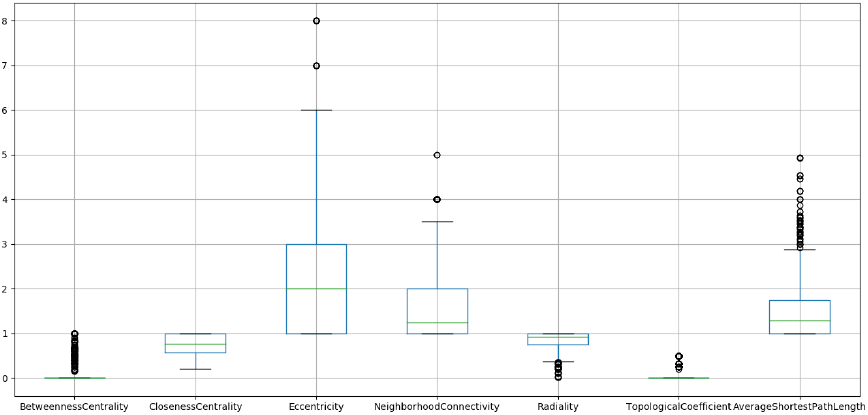
Boxplots

**Table 1:** Graph statistics

| Statistic | Value |
| --- | --- |
| # of nodes | 646 |
| # of edges | 533 |
| Average # of neighbors | 1875 |
| Diameter | 8 |
| Radius | 7 |
| Characteristic path length | 3767 |
| Density | 0.125 |
| Heterogeneity | 0.622 |
| Centralization | 0.162 |

### Scientific basis for new associations

In order to verify the scientific basis of our new associations, we have searched on *PubMed*, the most important *Library of Medicine*, our top prediction of drug and disease connection. *PubMed* is a database with noumerous abstract illustrating articles of biomedical nature with the possibility of finding an external link referring to the whole article. It is yield by *National Center for Biotechnology Information (NCBI)*.

The Table 2 describes the new associations with a reference to an article exposing the evidence seen by other scientists.

**Table 2:**
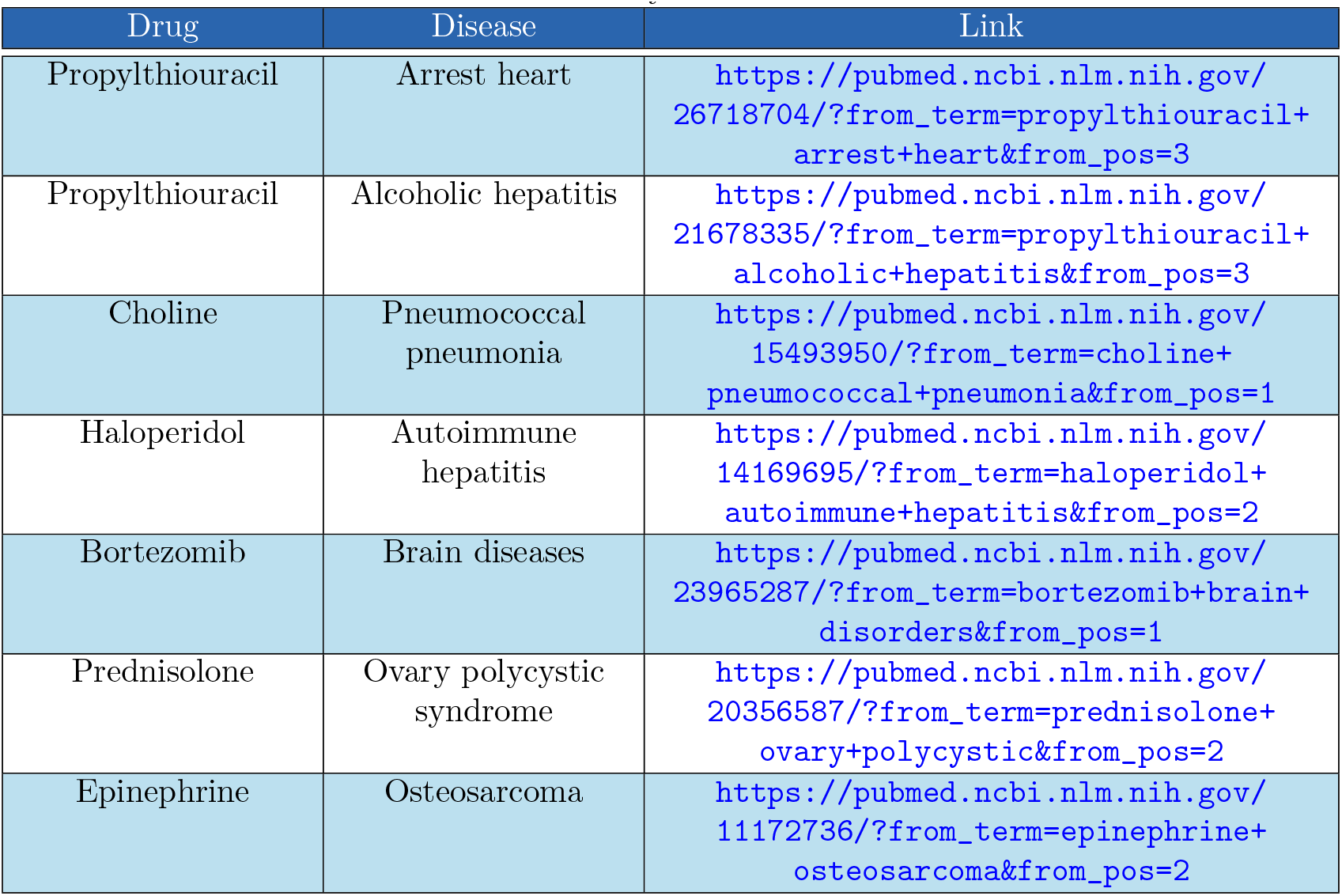
Summary of interaction data

| Drug | Disease | Link |
| --- | --- | --- |
| Propylthiouracil | Arrest heart | https://pubmed.ncbi.nlm.nih.gov/26718704/?from_term=propylthiouracil+arrest+heart&from_pos=3 |
| Propylthiouracil | Alcoholic hepatitis | https://pubmed.ncbi.nlm.nih.gov/21678335/?from_term=propylthiouracil+alcoholic+hepatitis&from_pos=3 |
| Choline | Pneumococcal pneumonia | https://pubmed.ncbi.nlm.nih.gov/15493950/?from_term=choline+pneumococcal+pneumonia&from_pos=1 |
| Haloperidol | Autoimmune hepatitis | https://pubmed.ncbi.nlm.nih.gov/14169695/?from_term=haloperidol+autoimmune+hepatitis&from_pos=2 |
| Bortezomib | Brain diseases | https://pubmed.ncbi.nlm.nih.gov/23965287/?from_term=bortezomib+brain+disorders&from_pos=1 |
| Prednisolone | Ovary polycystic syndrome | https://pubmed.ncbi.nlm.nih.gov/20356587/?from_term=prednisolone+ovary+polycystic&from_pos=2 |
| Epinephrine | Osteosarcoma | https://pubmed.ncbi.nlm.nih.gov/11172736/?from_term=epinephrine+osteosarcoma&from_pos=2 |

## Conclusion

The aim of this project was to identify existing drugs for other indications: this is the main core of *Drug Repurposing*. We used a computational approach based on *Network Medicine* exploiting a *Random Walker* that runs on a heterogeneous network. In BiRW we have seen how we combined the results of Similarity measures and Matrix factorization in order to run it different times to attain a new insight of the problem. Our aim, in fact, was to use different types of data in order to identify different similarities which cannot be seen using only one type. In Evaluation we have seen that our results are of interest and they are supported by scientific basis since we have found new associations that have previously been studied.

Drug repurposing was fundamental during the Covid19 pandemic since a drug prescribed for rheumatoid arthritis was used for patients in critical conditions undergoing ICU treatment. This example makes us reflect on the importance of this new field.

